# Assessment of anti-inflammatory bioactivity of extracellular vesicles is susceptible to error via media component contamination

**DOI:** 10.1101/2022.08.25.505301

**Authors:** Stephanie M. Kronstadt, Lauren Hoorens Van Heyningen, Amaya Aranda, Steven M. Jay

**Author notes:** Corresponding author: Steven M. Jay, Ph.D., Fischell Department of Bioengineering, University of Maryland, 3116 A. James Clark Hall, College Park, MD, 20742, USA.

## Abstract

Extracellular vesicles (EVs) are widely implicated as novel diagnostic and therapeutic modalities for a wide range of diseases. Thus, optimization of EV biomanufacturing is of high interest. In the course of developing parameters for a HEK293T EV production platform, we examined the combinatorial effects of cell culture conditions (i.e., static vs dynamic) and isolation techniques (i.e., ultracentrifugation vs tangential flow filtration vs size-exclusion chromatography) on functional characteristics of HEK293T EVs, including anti-inflammatory bioactivity using a well-established LPS-stimulated mouse macrophage model. We unexpectedly found that, depending on culture condition and isolation strategy, HEK293T EVs appeared to significantly suppress the secretion of pro-inflammatory cytokines (i.e., IL-6, RANTES) in the stimulated mouse macrophages. Further examination revealed that these results were most likely due to fetal bovine serum (FBS) EV contamination in HEK293T EV preparations. Thus, future research assessing the anti-inflammatory effects of EVs should be designed to account for this phenomenon.

## Introduction

With the potential to mirror the functional effects of parental cells and shuttle bioactive cargoes intercellularly, extracellular vesicles (EVs) hold promise as early-stage biomarkers and therapeutic interventions for various diseases as well as proficient drug delivery vehicles [1, 2]. As such, much effort has been expended in exploring the production, isolation, and functional characterization of EVs resulting in various methodologies that differ vastly across individual experiments and phases of research (i.e., preclinical vs clinical) [3]. In terms of EV production, there has been a collective movement to implement dynamic and/or 3D cell culture microenvironments, as these techniques are often more scalable than traditional flask culture and can produce EVs that resemble those within the physiological niche [4]. Specifically, bioreactor-based culture allows for the implementation of physiological parameters, including flow-derived shear stress, and has been shown to significantly increase EV production across numerous studies [5-14]. Importantly, the shift from static to dynamic culture is also known to cause dramatic changes in the EV therapeutic profile [9, 12, 14-16]; in some cases elucidating EV effects that were not present when utilizing traditional flask culture [15]. This phenomenon of altered EV bioactivity can also be observed when implementing different EV isolation strategies such as tangential flow filtration (TFF), ultracentrifugation (UC), or size-exclusion chromatography (SEC) [16, 17]. Thus, optimizing EV production and isolation methods may be vital in creating an ideal EV formulation for a given application.

Despite the important role that culture conditions and isolation strategies may play, only a couple of studies have explored the combinatorial effects of these techniques on perceived EV bioactivity [16, 18]. In one such study, Haraszti and colleagues demonstrated that EVs from umbilical cord-derived MSCs cultured within a microcarrier-based bioreactor and isolated via TFF were more efficient in delivering functional therapeutic siRNA into neurons than those EVs isolated from the same bioreactor culture using UC. They also show that bioreactor EVs isolated using UC were no more effective in siRNA transfer than EVs from traditional flask culture isolated using UC or TFF [16]. Previous research also cautions over assuming that the EVs are solely responsible for therapeutic activity, as many isolation strategies can result in the co-isolation of process impurities [19]. Demonstrating this is a study by Whitaker et al. in which the co-isolated and non-EV associated VEGF within MSC EV preparations had pro-angiogenic and pro-migratory effects on endothelial cells that could wrongfully be accredited to the EVs [20]. Altogether, these studies emphasize the importance of optimizing both cell culture conditions and isolation methods simultaneously while implementing careful functional assessment to produce the most viable EV therapeutic platform.

As for functional evaluation of EVs, much interest is often paid to EV anti-inflammatory capacity, as inflammatory dysregulation is ubiquitous in many chronic diseases (e.g., sepsis, cancer, and autoimmune disorders) and tissue regeneration issues [21, 22]. Recently, Pacienza and colleagues developed an *in vitro* LPS-stimulated mouse macrophage assay to assess the anti-inflammatory potential of mesenchymal stem cell (MSC)-derived EVs. Notably, results obtained in this *in vitro* assay were able to predict the efficacy of different MSC EV preparations in suppressing LPS-stimulated inflammation in mice [23]. As such, this assay and derivatives of it have been used regularly to test the anti-inflammatory properties of EVs [24-26].

In the present investigation, we examined the combinatorial effects of cell culture conditions (i.e., static vs dynamic) and isolation techniques (i.e., UC vs TFF vs SEC) on the size, morphology, and functional characteristics of EVs from HEK293T cells – a highly scalable source for the production of therapeutic EVs [27]. Surprisingly, depending on culture condition and isolation strategy, HEK293T EVs appeared to exert immunosuppressive effects in an LPS-stimulated mouse macrophage model. We found that this result was most likely due to FBS EV contaminants within the HEK293T EV samples. Our results highlight the importance of recognizing and controlling for potential contaminants in EV preparations when utilizing this assay.

## Methods

### Cell Maintenance

Human embryonic kidney cells (HEK293T) were purchased from ATCC (CRL-3216) and cultured in T175 flasks using Dulbecco’s Modification of Eagle’s Medium (DMEM) [+] 4.5 g/L glucose, L-glutamine, and sodium pyruvate (Corning, 10-013-CV) supplemented with 10% fetal bovine serum (FBS; VWR, 89510-186) and 1% penicillin-streptomycin (P/S; Corning, 30-002-CI). All HEK293T cells used in experiments were between passages 4-7. RAW264.7 mouse macrophages (ATCC, TIB-71) were cultured in T175 flasks with DMEM [+] 4.5 g/L glucose, L-glutamine, and sodium pyruvate (Corning, 10-013-CV) supplemented with 5% FBS and 1% P/S. RAW264.7 at passages 10-13 were used in experiments. Human umbilical vein endothelial cells (HUVECs) pooled from multiple donors (PromoCell; C-12203) were cultured in T75 flasks coated with 0.1% gelatin at 37 ^°^C for 1 h prior to seeding. HUVECs were maintained in complete endothelial growth medium-2 (EGM2; PromoCell, C-22111) supplemented with 1% P/S. During experiments, HUVECs were cultured in endothelial basal medium-2 (EBM2; PromoCell, C-22221) supplemented with 0.1% FBS and 1% P/S. HUVECs at passages 3 and 4 were used in experiments.

### HEK293T Cell Culture Conditions

#### Flask and Scaffold Culture

As static controls, HEK293T cells were cultured within T75 tissue culture flasks (VWR, BD353136) or a 3D-printed scaffold. The scaffold was printed from a biocompatible acrylate-based material (E-Shell 300; EnvisionTEC) using an EnvisionTEC Perfactory 4 Mini Multilens stereolithography apparatus (EnvisionTEC, Inc., Dearborn, MI, USA). The 12 cm^3^ volume construct contained a growth surface area (50 cm^2^) with small pillars (1 mm in diameter) that were spaced 2.5 mm apart. This design allowed efficient cell removal and immunofluorescence imaging as well as the facilitation of nutrient and gas transport and a mechanism to control fluid parameters predictably. Following printing, scaffolds were submerged in 99% isopropanol (Pharmco-Aaper, Shelbyville, KY) for 5 min to remove excess resin. The scaffolds were then flushed with fresh 99% isopropanol until all excess resin was removed after which the scaffolds were dried with filtered air. Resin curing was accomplished using 2,000 flashes of broad-spectrum light (Otoflash, EnvisionTEC, Inc.). Scaffolds were then cleaned in 100% ethanol for >30 min to eliminate any remaining contaminants, after which they were placed in fresh 100% ethanol and sterilized in an ultraviolet sterilizer (Taylor Scientific, 17-1703) for 10 min. Rehydration was achieved by soaking the scaffolds in sterile serial dilutions of ethanol to 1X PBS (pH 7.4) for 5 minutes per step (75:25, 50:50, 25:75, 0:100 ethanol:PBS). Rehydrated scaffolds were placed in 100% sterile 1X PBS in at 4°C until use. When ready to use, scaffolds were coated with 3 µg/cm^2^ fibronectin in sterile water at 37^°^C for 30 min. Flasks and scaffolds were seeded at a density of 2,500 cell/cm^2^ in 15 mL of media.

#### Perfusion Bioreactor Culture

For the perfusion bioreactor condition, select seeded scaffolds were hooked up to a Masterflex L/S Digital Peristaltic Pump (Cole-Parmer; Vernon Hills, IL, USA) using platinum-cured silicon tubing (Cole-Parmer, EW-95802-04). The scaffolds were operated at a flow rate of 5 mL/min which corresponded to a shear stress value of 3×10^−3^ dyn/cm^2^ as determined previously by computational fluid modeling via the Flow Simulation add-in for SolidWorks (Dassault Systèmes, Velizy-Villacoublay, France). The set-up allowed the scaffolds to be connected to a media reservoir containing 50 mL of EV-depleted FBS media. The pump and scaffolds were then placed within a cell culture incubator at 37°C with 5% CO_2_.

### Cell Staining and Imaging

After 24 h culture in the bioreactor, the scaffolds were filled with 4% paraformaldehyde for 15 min at room temperature to fix the HEK293T cells. Scaffolds were then washed three times with 1X PBS and filled with permeabilization buffer (300 µM sucrose, 100 µM sodium chloride, 6 µM magnesium chloride, 20 µM HEPES, and 0.5% Triton-X-100 solution) for 5 min at room temperature. AlexaFluor 488 Phalloidin in PBS (1:100) was used to stain cell actin and cells were visualized on a Nikon Ti2 Microscope (Nikon, Minato City, Tokyo, Japan).

### Media EV-Depletion Protocols

To produce EV-depleted FBS, heat-inactivated FBS was centrifuged at 118,000 × *g* for 16 h and the resulting supernatant was filtered through a 0.2 μm bottle top filter for use in subsequent media supplementation. To produce EV-depleted whole media, DMEM with full supplementation (10% heat-inactivated FBS and 1% P/S) was centrifuged at 118,000 × *g* for 16 h and the supernatant was sterile filtered (0.2 μm) for direct use in ensuing experiments.

### EV Isolation Techniques

During experiments, HEK293T culture media was replaced with media made with 10% EV-depleted FBS and 1% P/S (i.e., EV-depleted FBS media) or EV-depleted whole media. Approximately 100 mL of HEK293T conditioned media from each culture condition (i.e., flask, scaffold, or bioreactor) was subjected to a series of differential centrifugation to clear the media of cells as previously described [28]. In brief, conditioned media was first centrifuged at 1,000 × *g* for 10 min to remove any cells that may have detached during collection. The supernatant was then collected and centrifuged at 2,000 × *g* for 20 min to remove any larger cellular debris. This supernatant was then centrifuged at 10,000 × *g* for 30 min to remove any large organelles remaining. The cleared conditioned media was then subjected to the various isolation techniques described below. Final EV samples were stored at 4°C and were analyzed within three days of collection.

#### Ultracentrifugation

For ultracentrifugation (UC), the conditioned media was subjected to an additional centrifugation step of 118,000 × *g* for 2 h in a Type 70 Ti ultracentrifuge rotor (Beckman Coulter). The resulting EV pellet was resuspended in 1X PBS and placed into a Nanosep 300 kDa MWCO spin column (Pall, OD300C35) and spun at 8,000 × *g* until all PBS filtered through the membrane (∼10-12 min). EVs were washed two more times with 1X PBS, resuspended in 1X PBS, and sterile filtered using a 0.2 μm syringe filter.

#### Tangential Flow Filtration

Tangential flow filtration (TFF) was performed using a KrosFlo KR2i TFF system (Spectrum Labs, Los Angeles, CA, USA) equipped with a 100 kDa MWCO hollow fiber filter comprised of a modified polyethersulfone membrane (Spectrum Labs, D02-E100-05-N). Prior to processing the cleared conditioned media, the filter was first washed with at least three volumes of 1X PBS to remove the bacteriostatic reagent. Each filter was used no more than five times. To keep a shear rate at 4000 s^-1^, the flow rate was kept constant at 106 mL/min. All samples were processed at a transmembrane pressure (TMP) of 5 psi. Samples were first concentrated to a volume of 25 mL and then diafiltrated five times in 1X PBS to exchange buffers. Following buffer exchange, the EVs were concentrated to a volume of 6-9 mL. Further concentration was performed using a 100 MWCO centrifugal concentrator (Corning, 431486) to achieve a final volume of ∼0.5 mL. The final EV suspension was sterile filtered using a 0.2 μm syringe filter.

#### Size-Exclusion Chromatography

EVs were isolated via size-exclusion chromatography (SEC) using qEV Original columns (Izon Science, ICO-35) per the manufacturer’s protocol. Briefly, the cleared conditioned media was concentrated to 0.5 mL using the TFF set-up and centrifugal concentrators described in the previous section. After flushing the columns with 1X PBS, 500 μl of the concentrated media was applied to the top of the column and the first four 0.5 mL fractions after the void volume were collected and pooled. The pooled fractions were then concentrated using 100 MWCO centrifugal concentrators to 0.5 mL and sterile filtered (0.2 μm).

### Media Testing

To assess the effects of media components, various media formulations were subjected to the aforementioned isolation techniques. Unconditioned media test formulations (i.e., media without exposure to cells) were defined as the following: no cell (media without any cell conditioning), + FBS (media supplemented with EV-depleted FBS), or -FBS (media without FBS). To evaluate the effects of EV-depletion protocols used to remove contaminating EVs from the FBS components, HEK293T cells cultured in flasks were exposed to either EV-depleted FBS media or EV-depleted whole media as previously described (see Media EV-depletion Protocols section).

### EV Characterization

#### Protein and Particle Quantification

Total protein concentration was determined using BCA methods (G-Biosciences, 786-571) and size distribution as well as particle concentration were assessed using a NanoSight LM10 (Malvern Instruments; Malvern, UK) with Nanoparticle Tracking Analysis (NTA) software version 2.3. For each sample, three 30-second videos were captured with a camera level set at 12. EV samples were diluted to obtain 20-100 particles per frame and at least 200 completed tracks per video to ensure accurate analysis. The detection threshold was set and kept constant across all replicates and samples. The total number of EVs was evaluated using the final resuspension volume and then divided by the number of cells to give final data expressed as total number of EVs per cell.

#### EV Markers

Western blot analysis was used to determine the presence of specific EV markers as well as the purity of each sample. Based on the BCA results, 7 μg of protein from each EV sample was used for analysis and compared with 7 μg of cell lysate. EV markers were assessed using primary antibodies for Alix (Abcam, ab186429), TSG101 (Abcam, ab125011), and CD63 (Proteintech, 25682-1-AP), while the absence of contaminating proteins were confirmed using antibodies for GAPDH (Cell Signaling Technology, 2118) and calnexin (Cell Signaling Technology, 2679). All primary antibodies were added at a 1:1,000 dilution, excluding GAPDH which was diluted 1:2,000. A 1:10,000 dilution of a goat anti-rabbit secondary (LI-COR Biosciences, 926-32211) was used. Protein bands were imaged using a LI-COR Odyssey CLX Imager and analyzed using the associated software.

#### Transmission Electron Microscopy

EV morphology was visualized via transmission electron microscopy (TEM) using a negative staining technique. A portion of each EV sample (10 μl) was fixed in a 1:1 solution using 4% EM-grade paraformaldehyde (Electron Microscopy Sciences, 157-4-100) for 30 min at room temperature. A 10 μl droplet of the EV-PFA mixture was then allowed to adsorb to a carbon-coated copper grid (Electron Microscopy Sciences, CF200-Cu-25) for 20 min. After a brief wash using a of a drop of 1X PBS, the EV-coated grid was then placed on a drop of 1% glutaraldehyde (in 1X PBS) for 5 min. The grid was washed 5-7 times (2 min each wash) on deionized water droplets with blotting on filter paper between washes. The grid was then positioned on a droplet of uranyl-acetate replacement stain (Electron Microscopy Sciences, 22405) and allowed to dry completely for 10 min. Once prepared, the grids were imaged at 200 kV on a JEOL JEM 2100 LaB6 TEM.

### Bioactivity Assessment

#### Macrophage Stimulation Assay

To assess the effects of EV isolation technique and culture method on downstream applications, a mouse macrophage stimulation assay was utilized [23, 29]. RAW264.7 mouse macrophages were seeded at 60,000 cells/well in triplicate in a 48-well plate. 24 h later, two groups (i.e., six wells) of the macrophages were pre-treated with just media with PBS (vehicle control). Another group was pretreated using media supplemented with dexamethasone at 1 μg/mL (resuspended in 1X PBS) which served as the positive control (Dex; Sigma-Aldrich, D4902-25 MG). The remaining macrophages were pre-treated with the HEK293T EVs (resuspended in 1X PBS) from the various culture and isolation techniques at 5E9 EVs/mL diluted in cell culture media. After 24 h of incubation, the pre-treatments were removed and the macrophages were washed once with sterile 1X PBS. Cells were then either treated with just media spiked with PBS (vehicle control) or media spiked with lipopolysaccharide at 10 ng/mL (LPS; resuspended in 1X PBS; Sigma-Aldrich, L4391-1MG) for 4 h. The supernatants were then collected from the RAW264.7 macrophages and frozen at -80° C. Levels of secreted IL-6, RANTES, and TNF-α were assessed using the appropriate DuoSet ELISA kit (R&D Systems, DY406, DY478, DY410).

#### Tube Formation Assay

HUVECs were used to evaluate effects on endothelial tube formation as previously described [30]. In brief, HUVECs were trypsinized, counted, and aliquoted into a tube marked for each treatment that was filled with 2 mL of EBM2 supplemented with 0.1% FBS and 1% P/S. The cells were then pelleted at 220 × *g* after which the supernatant was removed and the cells were aliquoted to a final concentration of 120,000 cells/mL into complete growth media (positive control), basal media (EBM2 plus 0.1% FBS and 1% P/S) devoid of EVs (negative control), or basal media supplemented with EVs (5E9 EVs/mL) and gently but thoroughly resuspended. The treatments were then applied in triplicate at 500 μl per well (i.e., 60,000 cells/well) in a 24-well plate coated with growth factor reduced Matrigel (Corning, 354230). Cells were imaged at 6 h using a Nikon Eclipse Ti2 Microscope and the number of loops formed by the HUVECs was quantified using ImageJ.

#### Gap Closure Assay

To assess endothelial migration, HUVECs were seeded at 15,000 cells/well in a 0.1% gelatin-coated 96-well plate and allowed to grow to confluency. The monolayer was then disrupted using an AutoScratch (BioTek Instruments; Winooski, VT, USA) to create a cell gap meant to simulate a wound. HUVECs were gently washed with 1X PBS and serum-starved for 2 h via incubation with basal media. Following serum starvation, the media was aspirated and replaced with complete growth media (positive control), basal media (negative control), or basal media spiked with EVs (5E9 EVs/mL). The cell gap was imaged at 0 h and 20 h using a Nikon Eclipse Ti2 Microscope and the change in the gap area was calculated using ImageJ as previously described [31].

### Statistics

Data are presented as mean ± standard error of the mean (SEM). Two-way ANOVAs with Tukey’s multiple comparisons tests were used to determine statistical differences (p < 0.05) among groups across cell culture conditions and EV isolation techniques in the *in vitro* stimulated macrophage assay. One-way ANOVAs with Tukey’s multiple comparisons tests were used to detect statistical differences (p < 0.05) among groups in the *in vitro* gap closure assay and tube formation assay. All statistical analyses were performed using Prism 9.1 (GraphPad Software, La Jolla, CA). Notation for significance in figures are as follows: ns – p > 0.05: *-p < 0.05; ** - p < 0.01; *** or ### - p < 0.001; **** or ####-p < 0.0001).

## Results

### Cell culture condition and EV isolation technique have no effect on EV size or morphology

As various factors including cell culture methods and isolation techniques have been shown to influence downstream EV efficacy [32], we set out to dissect the effect of these factors on HEK293T EVs. HEK293T cells were chosen based on their proven ability to be integrated into scalable biotech processes as well as demonstrated ability to be engineered for therapeutic EV production [27, 33]. The EVs that they secreted are also thought to be relatively therapeutically inert, especially when compared with the EVs of other cell types (e.g., mesenchymal stem cells; MSCs) [34-36]. Thus, analysis of HEK293T EVs in biological assays could reveal confounding factors in commonly used EV production and characterization strategies. In the current study, HEK293T cells were cultured in a 3D-printed, acrylate-based scaffold hooked up to a peristaltic pump and operated at a 5 mL/min flow rate (3×10^−3^ dyn/cm^2^), constituting a set-up which will be referred to as the bioreactor. Prior studies in our lab have revealed that this specific flow rate allows adequate cell viability and increases EV production (i.e., EVs per cell) [5]. HEK293T cells were also cultured in static scaffolds and T75 flasks as controls. Conditioned media from the various culture methods were collected and EVs were isolated via ultracentrifugation (UC), tangential flow filtration (TFF), or size exclusion chromatography (SEC) as depicted in **Figure 1A**. There were no significant alterations in EV mode size as measured by NTA (Flask UC – 96.9 ± 5.5 nm; Flask TFF – 86.3 ± 6.4 nm; Flask SEC – 90.9 ± 10.1 nm; Scaffold UC – 88.8 ± 6 nm; Scaffold TFF – 92.9 ± 3.4 nm; Scaffold SEC 90.9 ± 9.5 nm; Bioreactor UC – 98.7 ± 5.6 nm; Bioreactor TFF – 84.2 ± 4 nm; Bioreactor SEC 90.1 ± 2 nm) (**Figure 1B**). Additionally, greater than 90% of the EV population from each sample were well within the range of acceptable exosomal diameter (40 – 200 nm) (**Figure 1B**) (Andaloussi et al. 2013). Immunoblotting established the presence of specific EV markers (Alix, TSG101, and CD63) and the absence of cellular debris indicators (Calnexin and GAPDH) (**Figure 1C**). TEM images showed no effect of culture or isolation technique on HEK293T EV morphology (**Figure 1D**).

**Figure 1.**
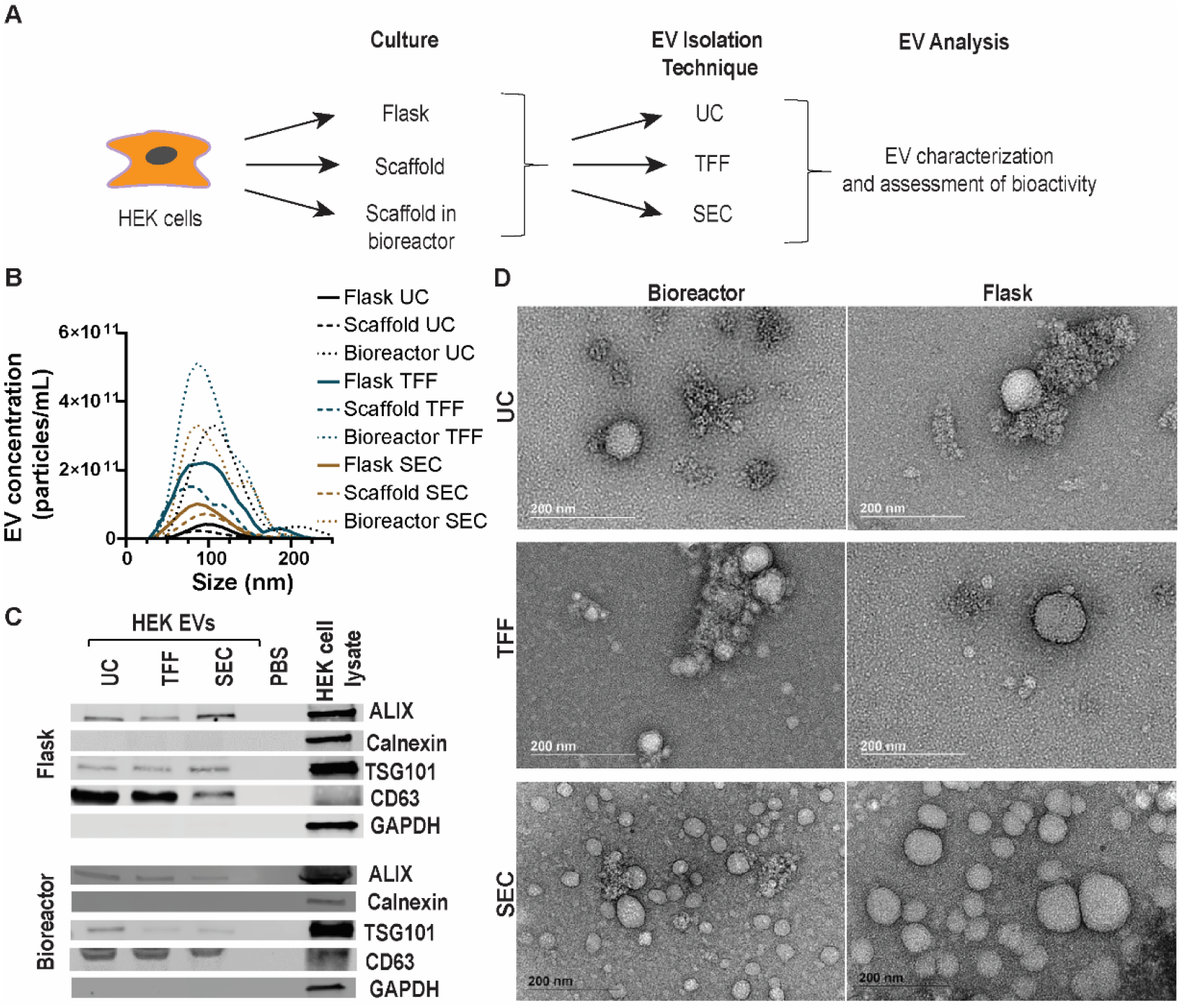
Culture conditions and isolation techniques do not impart significant effects on size or morphology of HEK293T EVs. (A) Schematic depicting the experimental workflow. (B) Size distribution of HEK293T-derived EVs generated and isolated via the various culture conditions and isolation methods (ultracentrifugation, UC; tangential flow filtration, TFF; size exclusion chromatography, SEC). (C) Immunoblotting of HEK293T EVs from flask and bioreactor conditions isolated via UC, TFF, or SEC for EV-specific markers (CD63, Alix, TSG101) and cell markers (Calnexin and GAPDH). (D) TEM images of HEK293T EVs isolated from the flask and bioreactor culture conditions using each isolation method. Data and images are representative of two independent experiments (N = 2).

#### Immunomodulatory activity of HEK293T EV preparations is altered by culture condition and isolation method

HEK293T EV isolates prepared using the various culture conditions (i.e., flask, scaffold, or bioreactor) and separation methods (i.e., UC, TFF, or SEC) were assessed in an LPS-stimulated mouse macrophage assay, with final read-outs being the secretion levels of several inflammatory cytokines (i.e., IL-6, RANTES, TNF-α). These cytokines are known to regulate inflammation and are correlated with EV immunomodulatory activity *in vivo* [23, 37, 38]. It was observed that levels of the secreted inflammatory cytokines differed significantly when macrophages were exposed to HEK293T EV preparations from the various culture methods and isolation technique combinations (**Figure 2**). For every cytokine analyzed, there was a significant interaction of culture technique with isolation method as determined by a two-way ANOVA (IL-6: p < 0.0001; RANTES: p < 0.0001; TNF-α: p = 0.0056). Surprisingly, Tukey’s multiple comparison tests revealed that HEK293T EV preparations from every condition, except those from the bioreactor culture isolated using SEC, significantly reduced the levels of IL-6 secretion compared with the LPS-only group (**Figure 2A**). A similar trend, although not always significant, was observed in the levels of secreted RANTES (**Figure 2B**). Isolating via the UC method in particular produced preparations with a significantly increased ability to suppress the secretion of inflammatory cytokines (**Figure 2**); an effect that was often lost if cells were cultured within the bioreactor (**Figure 2B, C**). Notably, preparing EV isolates via TFF or SEC weakened the observed suppression of cytokine secretion.

**Figure 2.**
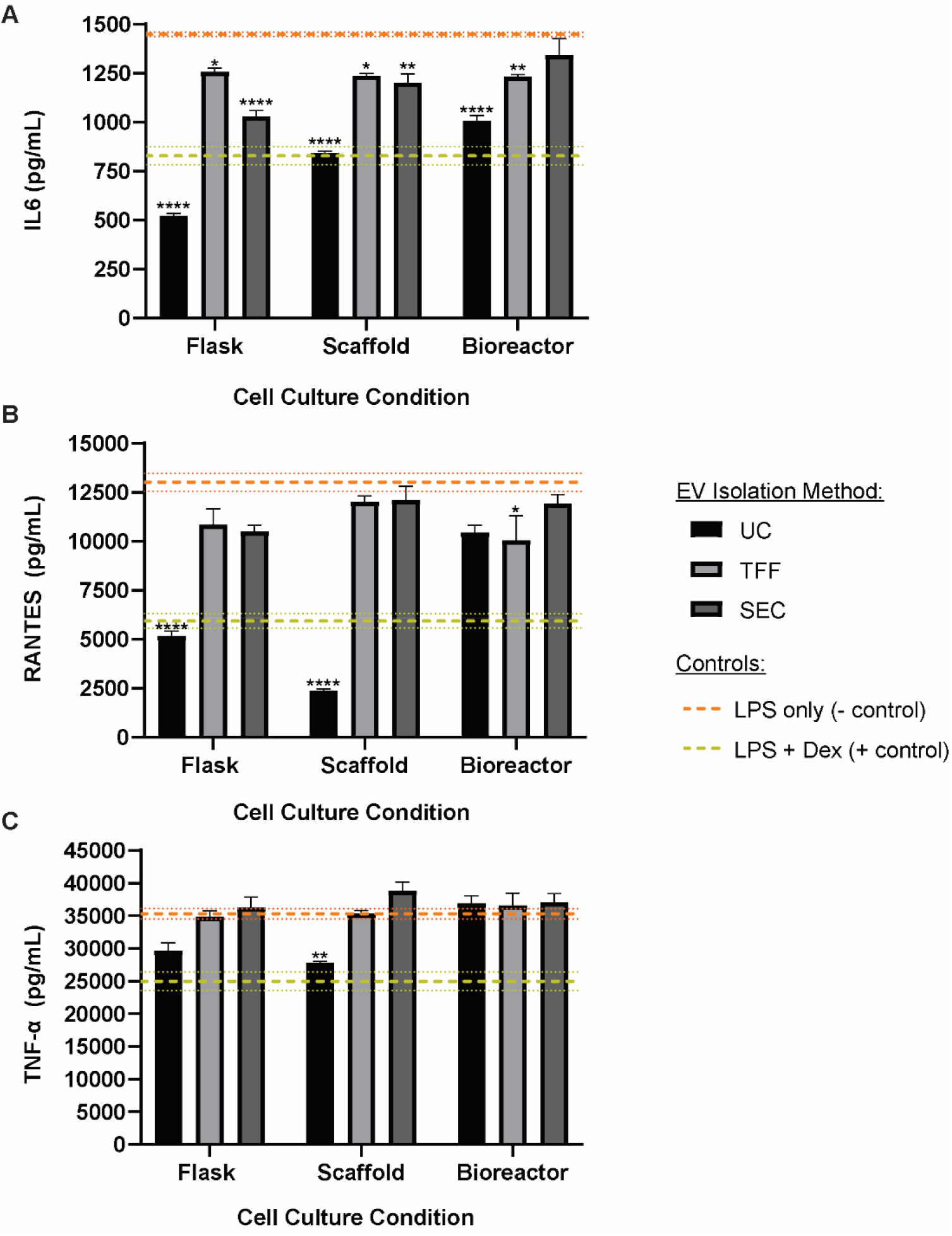
Culture conditions and isolation techniques alter the immunomodulatory activity of HEK293T EVs. RAW264.7 mouse macrophages were pretreated with either HEK293T EVs from cells within the various culture conditions (i.e., flask, scaffold, or perfusion bioreactor) and isolated using different techniques (i.e., ultracentrifuge – UC, tangential flow filtration – TFF, or size-exclusion chromatography – SEC) or with dexamethasone (Dex) prior to stimulation with lipopolysaccharide (LPS). RAW264.7 supernatants were collected and (A) IL-6, (B) RANTES, and (C) TNF-α secretion was quantified via ELISAs. Data is plotted as mean and error bars represent the standard error of the mean (SEM). Control means are represented as dashed lines with dotted lines representing the SEM. Data includes three technical replicates and data are representative of two independent biological replicates (N = 2). Statistical significance was calculated using a two-way ANOVA using Tukey’s multiple comparison tests (*-p < 0.05; ** - p < 0.01; **** - p < 0.0001 compared to LPS negative control).

#### Media components influence the results of the macrophage stimulation assay

To determine if media components affect the readout of the macrophage stimulation assay, unconditioned media (i.e., media not previously exposed to HEK293T cells) and conditioned media collected from HEK293T cells cultured in flasks were exposed to the various isolation techniques (i.e., UC, TFF, or SEC) and applied to RAW264.7 macrophages prior to LPS stimulation. Interestingly, both unconditioned and conditioned media preparations significantly and similarly reduced the secretion of IL-6 from the stimulated macrophages regardless of isolation technique (**Figure 3A**). Analysis of RANTES concentrations reveals a reduction of the cytokine only in the cells treated with the UC-isolated preparations, with a significant reduction in RANTES secretion when treated with the conditioned media (**Figure 3B**). However, when looking at TNF-α, the trends readily apparent in the IL-6 concentrations were lost, except for a reduction when isolating with TFF (**Figure 3C**).

**Figure 3.**
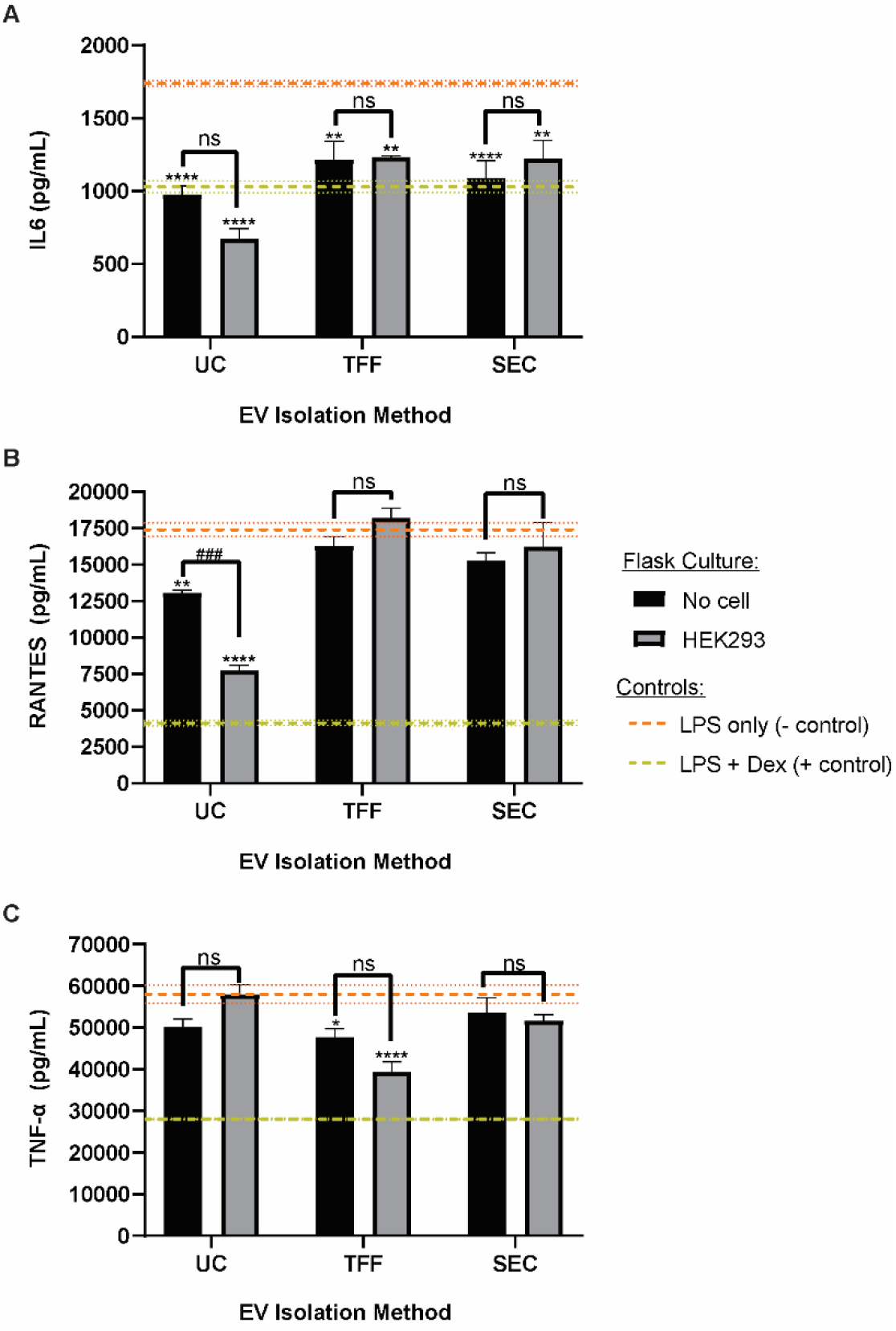
Unconditioned media subjected to each isolation technique alter cytokine secretion in stimulated macrophages in a similar manner to HEK293T EV preparations. Unconditioned media (i.e., no prior exposure to cells; ‘no cell’) or media conditioned with HEK293T cells within flasks were exposed to each isolation procedure (i.e., ultracentrifugation – UC, tangential flow filtration – TFF, or size-exclusion chromatography – SEC). Either the resulting preparations or dexamethasone (Dex) were applied to RAW264.7 mouse macrophages prior to stimulation with lipopolysaccharide (LPS). RAW264.7 supernatants were collected and (A) IL-6, (B) RANTES, and (C) TNF-α secretion was quantified via ELISAs. Data is plotted as mean and error bars represent the standard error of the mean (SEM). Control means are represented as dashed lines with dotted lines representing the SEM. Data includes three technical replicates and data are representative of two independent biological replicates (N = 2). Statistical significance was calculated using a two-way ANOVA using Tukey’s multiple comparison tests (ns – p > 0.05; *-p < 0.05; ** - p < 0.01; **** - p < 0.0001 compared to LPS negative control).

To determine if these results were confined to the stimulated macrophage assay, unconditioned media were isolated via UC and applied in two orthogonal *in vitro* assays. Results showed that in both an endothelial gap closure assay and an endothelial tube formation assay, unconditioned media did not induce any significant response and were similar to the negative control (i.e., basal media) (**Figure 4**).

**Figure 4.**
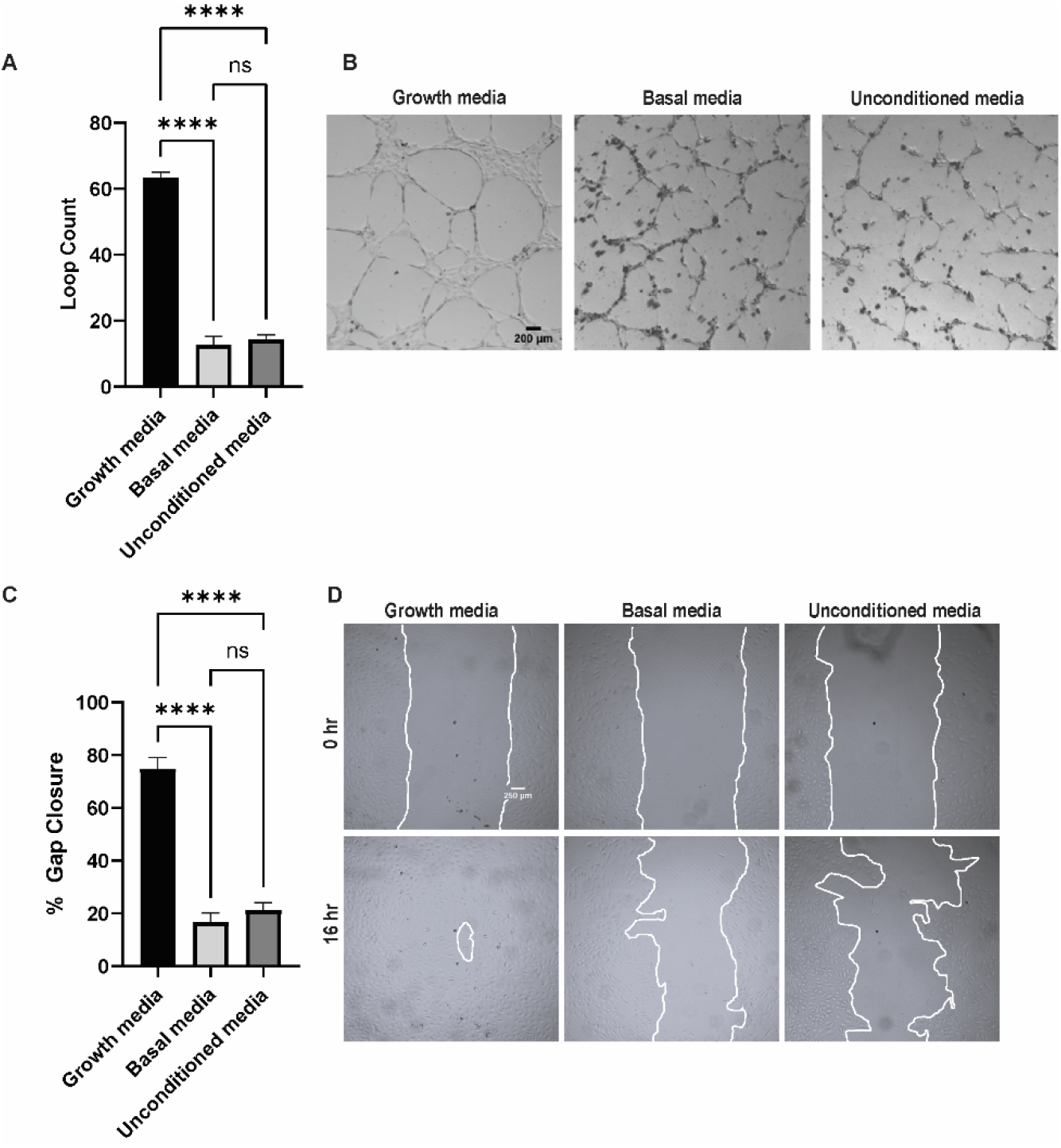
Unconditioned media do not elicit any significant response in *in vitro* angiogenic assays. (A) Number of loops formed by human umbilical vein endothelial cells (HUVECs) in a tube formation assay after treatment with growth media (positive control), basal media (negative control), or unconditioned media following exposure to the ultracentrifugation isolation protocol along with (B) the corresponding representative images at 16 hr. (C) Percent gap closure 16 h after treatment with growth media, basal media, or unconditioned media subjected to ultracentrifugation. There were no significant differences between the unconditioned media treatment and the negative control (basal media) in either assay (p > 0.05). All images and data are representative of two biological replicates with three technical replicates each. Statistical differences were analyzed using a one-way ANOVA with Tukey’s multiple comparisons test (ns – p > 0.05, **** - p < 0.0001).

To further dissect the observed effects, unconditioned media containing FBS (i.e., + FBS) or unconditioned media without FBS (i.e., - FBS) underwent the UC isolation protocol and were tested in the stimulated macrophage assay. The results revealed that the suppression of IL-6 secretion from macrophages was lost when FBS was absent from the media (**Figure 5A**). A similar trend was observed when looking at RANTES levels, but only when using the UC isolation method (**Figure 5B**). All patterns of cytokine suppression were diminished when examining TNF-α levels (**Figure 5C**). Based on these results, the EV-depletion protocols for media supplementation were examined. Two disparate protocols were followed, and the media tested again in the macrophage assay. In brief, FBS was either depleted of EVs prior to addition into the media (i.e., FBS-depleted) or after being added to the media (i.e., whole media-depleted), subjected to the UC isolation protocol, and applied to macrophages prior to LPS stimulation. Results showed that when whole media was depleted, the secretion of IL-6 from stimulated mouse macrophages was no longer subdued and was statistically similar to the LPS-only control (**Figure 6**).

**Figure 5.**
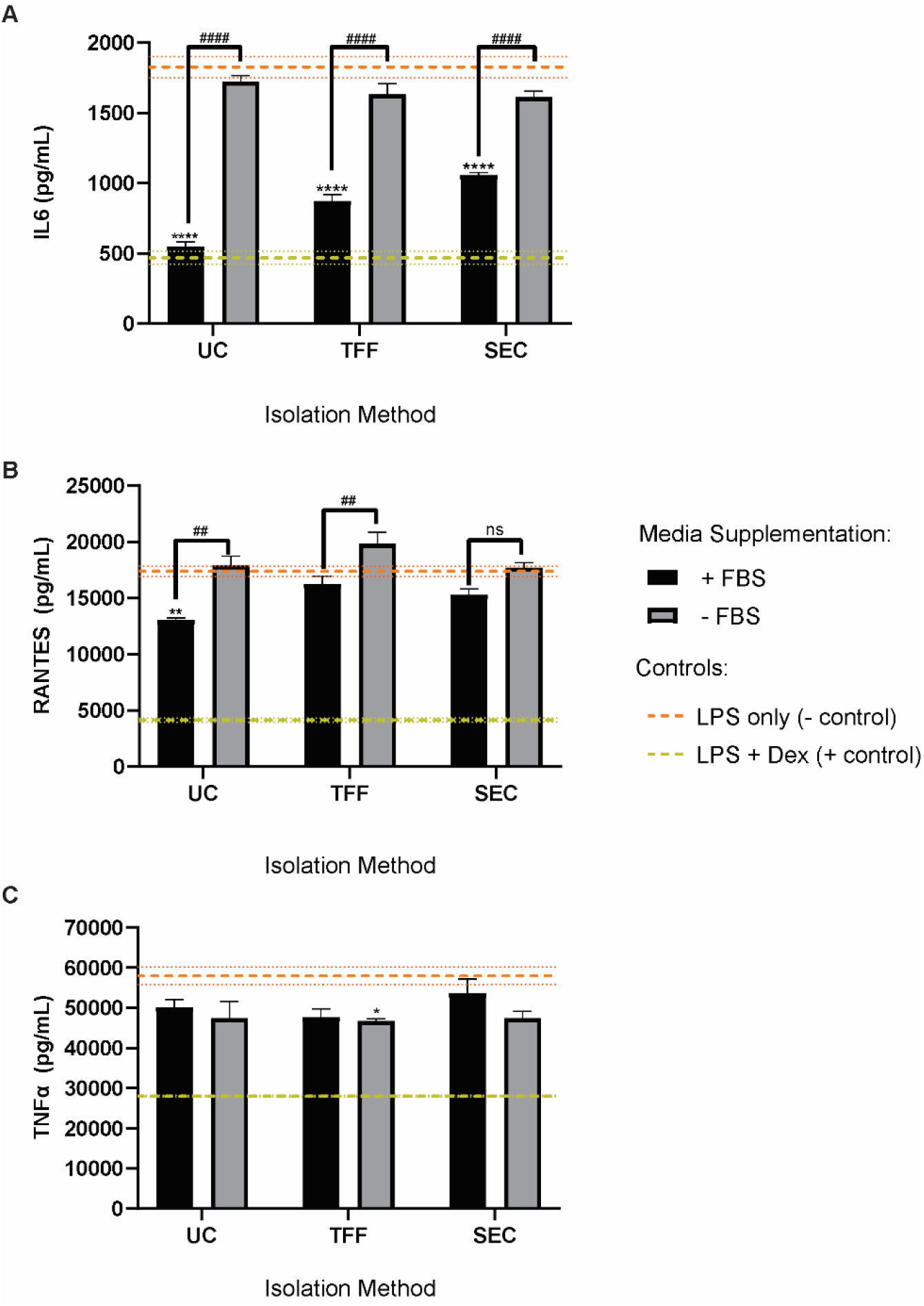
Fetal bovine serum (FBS) may be responsible for observed immunomodulatory responses of stimulated macrophages. Unconditioned media with FBS (+ FBS) and without FBS (-FBS) were subjected to a routine ultracentrifugation EV isolation protocol and applied to mouse macrophages prior to stimulation with LPS. The concentrations of (A) IL-6, (B) RANTES, and (C) TNF-α were quantified using ELISAs. Data is plotted as mean and error bars represent the standard error of the mean (SEM). Control means are represented as dashed lines with dotted lines representing the SEM. Data includes three technical replicates and data are representative of two independent biological replicates (N = 2). Statistical significance was calculated using a two-way ANOVA using Tukey’s multiple comparison tests (ns – p > 0.05; *- p < 0.05; ** - p < 0.01; **** - p < 0.0001 compared to LPS negative control; #### - p < 0.0001).

**Figure 6.**
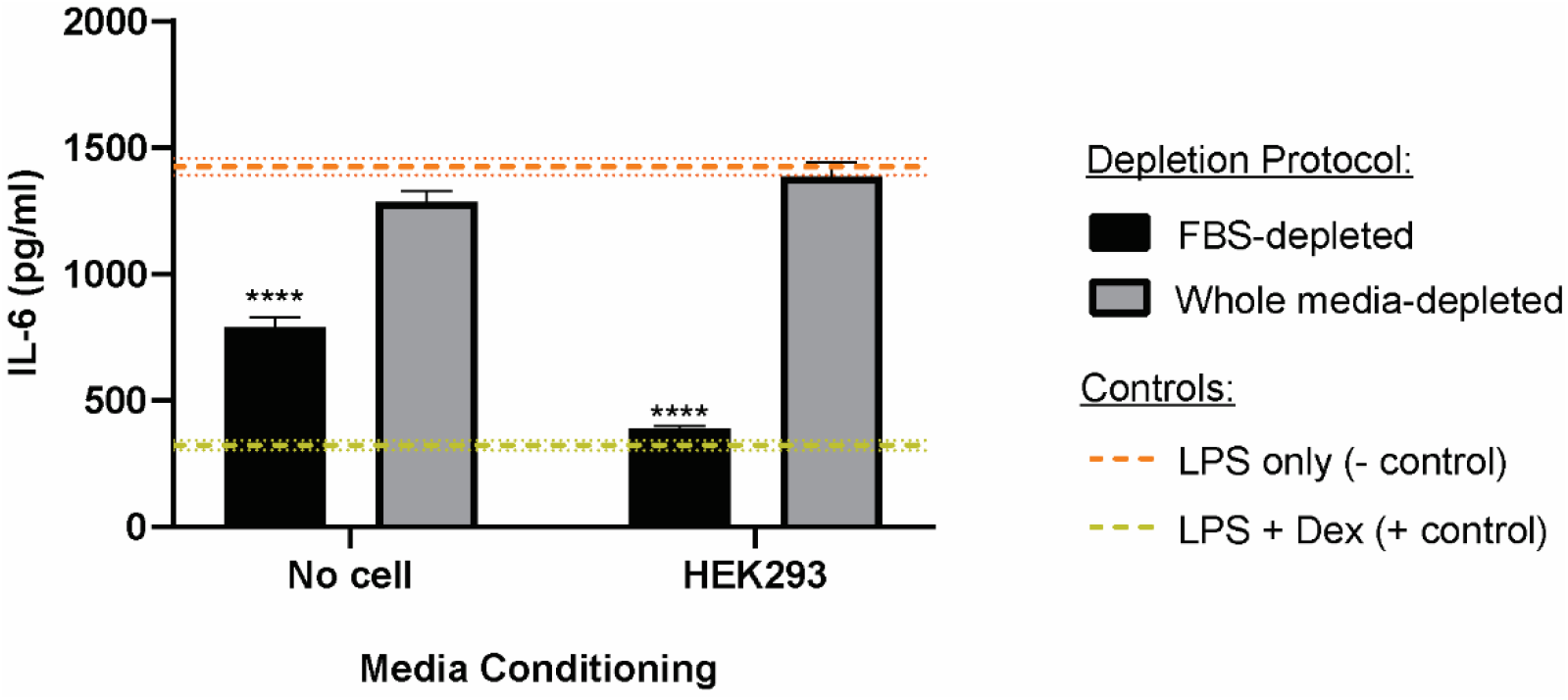
EV-depletion protocol of media supplements alter the results of the stimulated macrophage assay. FBS was either depleted of contaminating bovine EVs prior to addition to the media (FBS-depleted) or after addition (whole media-depletion), incubated either with (HEK293T) or without cells (no cell), subjected to the ultracentrifugation protocol, and applied to macrophages immediately prior to LPS. When the whole media protocol was used, IL-6 secretion was no longer suppressed. Data is plotted as mean and error bars represent the standard error of the mean (SEM). Control means are represented as dashed lines with dotted lines representing the SEM. Data includes three technical replicates and data are representative of two independent biological replicates (N = 2). Statistical significance was calculated using a two-way ANOVA using Tukey’s multiple comparison tests (**** - p < 0.0001 compared to LPS negative control).

## Discussion

It is well-known that cell culture parameters (e.g., cell density, passage, media composition, substrate architecture) as well as isolation strategies can significantly impact the molecular composition and therapeutic function of EVs [16, 17, 31, 39-49]. Particularly, this has important downstream implications as movement from the bench to the clinic requires implementing scalable GMP-compliant manufacturing techniques in both EV production (e.g., serum-free media and dynamic bioreactors) and EV isolation (e.g., size-exclusion chromatography), which are often quite different from those utilized in the preclinical setting and can be ever-changing even through clinical phases [3]. In the present study, we set out to investigate how these various upstream and downstream parameters may interact to alter EV characteristics using HEK293T cells as the EV source based on the proven compatibility of these cells with industry-scale bioprocess operations. To examine upstream parameters, we implemented a 3D-printed scaffold-perfusion bioreactor system previously utilized in our lab and exposed HEK293T cells to low levels of flow-derived shear stress (i.e., 3×10^−3^ dyn/cm^2^). To examine downstream isolation strategies, we isolated the subsequently secreted HEK293T EVs using the gold-standard technique of ultracentrifugation (UC) and compared it with the more scalable methods of size-exclusion chromatography (SEC) and tangential flow filtration (TFF). Our results suggest that EV size and morphology are not significantly altered when these particular cell culture parameters and isolation strategies are varied (**Figure 1**). It is important to evaluate both EV size and shape as changes in these characteristics are used as indicators of EV membrane integrity and have been known to be modified by external forces, particularly the shearing involved in high-speed centrifugation [17, 40]. When coupled with membrane markers, size can also be used to suggest possible EV biogenesis routes, as smaller EVs are thought to be derived more so from the endosomal compartment rather than the plasma membrane [50]. Previous research has shown similar results in which EV size and shape are not changed by the culture or isolation methodology used. For example, when comparing UC and TFF isolation methods following traditional flask and 3D microcarrier bioreactor-based culture of umbilical cord-derived MSCs, Haraszti and colleagues found all EV preparations to have similar size distributions [16]. Additionally, prior work in our lab has demonstrated no significant effect of dynamic culture on such physical parameters of EVs secreted from human dermal microvascular endothelial cells (HDMECs) [5]. However, a study by Sharma et al. showed significant differences in size distributions as well as nanoscale topography (i.e., surface roughness) that was evident across various breast cancer cell-derived EVs that were isolated using four different isolation methods (i.e., ultracentrifugation, density ultracentrifugation, immunoaffinity, precipitation) [51]. The disagreement in results could be attributed to differences in assessment techniques. While the aforementioned studies relied on nanoparticle tracking analysis and transmission electron microscopy, Sharma and colleagues used disparate methods to quantify and assess EV characteristics including atomic force microscopy (AFM), multi-angle light scattering (MALS), direct stochastic optical reconstruction microscopy (dSTORM), and microfluidic resistive pore sizing (MRPS). They also measured surface nano-roughness; a dimension rarely evaluated for EVs [52]. Importantly, surface roughness of synthetic nanoparticles has been proven to change the protein corona and subsequently alter their cellular uptake [53]. Although of biological origin, it makes sense that alterations in the surface roughness of EVs may impart similar changes in recipient cell internalization. Altogether these results suggest that, when possible, it is important to utilize orthogonal methods and consider unique characteristics to more rigorously assess EV biophysical parameters to help understand the mechanisms underlying downstream therapeutic effects.

We then assessed whether the culture or isolation techniques alter the bioactivity of HEK293T EVs by applying them in an *in vitro* LPS-stimulated mouse macrophage model. We found that the EVs, particularly those isolated using ultracentrifugation, significantly reduced the secretion of pro-inflammatory cytokines (i.e., IL-6 and RANTES); an effect that was largely lost when culturing within the bioreactor or isolating using TFF or SEC (**Figure 2**). As HEK293T EVs are not known to have significant immunomodulatory effects [34], these results were unexpected. Nevertheless, ultracentrifugation is notorious for the co-isolation of proteins and other small molecules which may be imparting the observed therapeutic effect [51, 54-56]. Moreover, previous research in our lab using a similar bioreactor has shown a significant reduction in total protein content per EV [5], which could help explain the loss of the observed effect when using EVs from HEK293T cells cultured within the bioreactor in the present study (**Figure 2**). This change in protein content in dynamic culture is to be expected as it has been shown that flow can alter the protein corona that forms on nanoparticles [57]. The composition of the EV protein corona under dynamic conditions should be evaluated in future studies.

To confirm whether the HEK293T EVs themselves were truly suppressing cytokine secretion, we focused only on traditional flask culture and assessed the effect of unconditioned media (i.e., media incubated without cells) after being processed using UC, TFF, and SEC. We found that the unconditioned media significantly reduced the IL-6 concentration regardless of isolation technique and lessened the RANTES and TNF-α concentrations particularly when isolating using UC and TFF, respectively (**Figure 3**). We found that this observation of the unconditioned media having an effect was only present in this specific assay, as there were no significant effects of unconditioned media subjected to ultracentrifugation in other *in vitro* assays commonly used to assess EV bioactivity (i.e., gap closure and tube formation) (**Figure 4**). This suggests an interaction between the component(s) of the media and the LPS response in this particular assay. Indeed, fetal bovine serum (FBS), a common reagent for media supplementation, has been shown to alter LPS sensitivity in cell cultures [58] as well alter EV analyses [59]. Thus, we tested unconditioned media without FBS supplementation and found that nearly all treatments were comparable to the LPS control, thus implicating FBS as the component responsible for the cytokine suppression (**Figure 5**). Interestingly, a previous study by Beninson and Fleshner found that FBS-derived EVs can have an immunosuppressive effect on primary rat macrophages [60]. Inspired by this, we revisited our EV-depletion protocol and found that differences in the methodology used to EV-deplete media can result in drastic differences in cytokine secretion (**Figure 6**). In the pursuit of understanding the effects of HEK293T EVs in the stimulated macrophage model, we were able to optimize our existing EV-depletion protocol which is a foundational component of the majority of our research.

As EV-based therapies have shown promise in a wide array of applications, there is much interest in the movement of these therapeutics into the clinic. However, successful clinical translation hinges upon the ability to optimize EV therapeutic effects within a scalable, GMP-compliant environment. In our work, we demonstrate an unexpected immunosuppressive effect of HEK293T EVs that differed depending on culture conditions of the HEK293T cells as well as isolation strategies used for the EVs. This may explain the disparity between studies that report no effect of unmodified HEK293T EVs in their bioactivity assays [61, 62] and others that describe significant immunomodulatory activity [63]. As some culture conditions and isolation methods may be more amenable to scale-up processes than others, it is important to consider these parameters as early as possible to ensure a smooth transition to clinical exploration. We also show that media components from the cell culture may contaminate the separated formulation and alter final EV therapeutic evaluation; a result that further emphasizes the urgency to move to xeno-free, chemically-defined culture conditions and to improve or introduce new downstream separation methods. As identity and purity are major quality considerations to move investigative new drugs into the realm of approved therapeutics [3], this work reiterates the necessity for multiple orthogonal assays as well as the consideration of all controls in order to confirm that any observed effects are truly the result of the experimental EV condition.

